# Expanding the use of reindeer foetal bone measurements for zooarchaeological applications

**DOI:** 10.1101/2024.03.28.587213

**Authors:** Emmanuel Discamps, Marie-Cécile Soulier

## Abstract

When foetal bones are preserved in archaeological sites, they are often used to identify the seasonality of prey acquisition by past human populations and, subsequently, to discuss their lifestyle, their management of food resources, nomadic cycles, etc. To do so, zooarchaeologists use charts to estimate foetal age based on the growth of their bones. For reindeer (*Rangifer tarandus*), a species that was widely exploited since the Palaeolithic throughout Eurasia, existing reference data are limited and require the measurement of complete bones, a procedure that is rarely applicable to archaeological contexts in which bones are often fragmented. In this study we present a wide range of measurements (9-10 measurements per bone) taken on the humerus, radius, metacarpal, femur, tibia and metatarsal of 31 individuals housed at the Zoological Museum of the University of Oulu (Finland). With this large data set, a more accurate estimation of the time of death of reindeer foetus can be achieved using skeletal measurements, even in the case of fragmented bones. To facilitate the use of this referential, an open-access web interface (<u>foetusmeteR</u>) was designed in RShiny. This interface allows for the direct estimation of foetal age and season of death by entering a single skeletal measurement, as well as the possibility of estimating if two bones might correspond to the same individual using two different measurements. This new tool should help to discuss in more detail the condition of reindeer herds hunted in the past, the hunting techniques and strategies that may have been used by human groups, and allow for a more detailed reconstruction of the seasonal nomadic cycle of past societies that focused their subsistence on *Rangifer* populations.

## Introduction

Apart from its key role in many modern Arctic herding societies, reindeer (*Rangifer tarandus*) was also one of the most commonly hunted prey throughout the Paleolithic in Eurasia, as testified by the abundance of its skeletal remains in archaeological sites (e.g. Delpech, 1983; Discamps et al., 2011; Boyle, 2017; Yaworsky et al., 2023). When foetal bones are found amongst archaeological assemblages, they constitute a valuable asset because these skeletal elements attest to the exploitation of pregnant females by past humans and allow for an estimation of the slaughter season of these reindeer females. By combining such information with other data, it then becomes possible to reconstruct the hunting strategies and techniques employed by past societies. The presence of foetal bones can however raise questions for scholars interested in the study of past human-reindeer interactions (i.e. zooarchaeologists).

The literature available on societies whose economy is still (or was until recently) mainly based on hunting provides some interesting food for thought on the subject of foetuses. First of all, their presence is to be expected on kill sites, where game is usually butchered first in order to remove unwanted parts and save weight for transport to base camps. Past human populations sometimes targeted specific social groups for precise needs, and foetus bones can testify of the hunt of matriarchal groups in order to obtain young individuals with very valuable hides. Foetal bones are however also found in “secondary” sites such as base camps, where they might have been introduced deliberately. The very tender foetal meat is generally given to people who have difficulty chewing, such as young children or the elderly (Jenness, 1922; Speck, 1935; Holston, 1963; Gubser, 1965; Burch, 1972; Hungry Wolf, 1980; Ingstad, 1992), and can be eaten boiled or roasted. Foetal skin is also highly prized for its fineness and can be used to make children’s clothing, blankets or fine summer clothes, masks and containers (e.g. Stefánsson, 1914; Curtis, 1930; Banfield, 1977; Binford, 1978, 1981; Grønnow et al., 1983; Russell, 1995; Parget, 2004; Pinson, 2004; Blackman, 2006). When pregnant females are hunted, some groups (such as the Quechua), include the foetus in offerings (Froemming, 2006). All of these uses, and certainly many others, are likely to date back to prehistoric times and can provide interesting food for thought about the presence of foetuses in archaeological sites.

When *Rangifer* foetal bones are found on an archaeological site, zooarchaeological studies often take advantage of them to try to access precious seasonal data (e.g. Castel, 1999; Morin, 2004; Paletta, 2005; Castel & Chauvière, 2007; Bemilli & Bayle, 2009; Magniez, 2009; Connet et al., 2012; Soulier, 2013, 2014; Pétillon et al., 2015; Castel et al., 2017; Fontana, 2017; Costamagno, 2021). To do so, such studies referred to the few published datasets on foetal bone growth for reindeer/caribou. To our knowledge, only two works carried out measurements of *Rangifer* foetal bones:

● Based on hindfoot length data from 4 individuals supplied by Kelsall (1957), Spiess (1979) noted a straight-line regression. Spiess proceeded to conclude that *“assuming a relatively constant growth rate and a relatively constant proportion between total hind-foot length and diaphyseal lengths of fore and hind-limb long bones, […] simple regressions [can be used by zooarchaeologists] to go from the diaphyseal length of a given fetal long bone, to total hind-foot length, [and thus] to fetal age”* (Spiess, 1979: pp. 94-95). He then compared hint-foot lengths and diaphyseal lengths on 6 young individuals to establish correlations. From this, he developed a simple formula, based on the total length of the complete bone, to estimate foetal development from the metapodials, femur, humerus and tibia. Zooarchaeologists have since mostly used these equations for seasonal estimation (cf. references cited above).
● Roine and collaborators (1982) studied the development of 348 reindeer foetuses from Finland, providing mean measurements of the metacarpal length per estimated age. They demonstrated strong correlations between foetus age, body weight and metacarpal length. Using the equation they provide, zooarchaeologists can estimate foetal age using a length measurement of a complete metacarpal bone.

These two studies are undoubtedly of capital importance for estimating the slaughter seasons of pregnant females. They do however suffer from three major drawbacks when it comes to their use for zooarchaeological analysis. First and foremost, these two studies were not designed to work on fragmented bones, even though foetal bones are highly prone to alteration by taphonomic processes due to their fragility and low density. To compensate for this bias, one solution is to estimate the completeness of the bone fragment (e.g. Soulier, 2013). However, a few millimetres of difference can alter our estimates by several weeks, or even months, which is not without consequences for discussions on the hunting techniques and strategies of prehistoric societies (depending on the season of death, zooarchaeologists would infer either hunting on isolated individuals or mass slaughters of migrating large herds). Secondly, no data is available for the radius. Given that foetal bones are not very dense and prone to destruction by taphonomic processes, the absence of one skeletal element is highly regrettable: important seasonal information cannot be accessed using radii, despite the fact that this precious and fragile material has been preserved over time. Finally, the lack of data for complete individuals in these studies (e.g. measurements of the humerus and femur of the same individual) makes it difficult to determine with a high degree of precision whether different skeletal elements could belong to the same or multiple individuals. This has implications for zooarchaeological interpretations of past hunting strategies, as well as considerations relating to the presence of foetuses in archaeological sites.

The objectives of our study are manifold:

- Increase the available reference measurements (currently limited, for most elements except the metacarpal, to Spiess’ estimates based on the 4 individuals measured by Kelsall), in order to provide more robust correlations;
- Provide measurements on isolated elements (i.e. the bones themselves) and not from hindfoot lengths, which are *de facto* estimates;
- Obtain seasonality data on all long bones, so as to mobilise the largest possible corpus for estimating the game killing season;
- Propose easily usable measurements on fragmented material by multiplying measurement points;
- Provide correlations between elements for accurate calculation of foetal Minimum Number of Individuals (MNI) when bones are found isolated, i.e. the fewest possible number of animals in a skeletal assemblage;
- Design an easy-to-use interface for the analysis of foetal measurements.

To these ends, a metric analysis of 31 *Rangifer tarandus* foetal bones is presented here.

## Material and methods

A total of 31 individuals stored at the Zoological Museum of the University of Oulu (Finland) were measured: 24 domesticated *Rangifer tarandus tarandus* (6 females, 11 males, 7 unknown), 6 wild Finnish forest reindeer *Rangifer tarandus fennicus* (3 females, 3 males) and one hybrid *R. t. tarandus-fennicus* female. They were all collected in Finland (in the municipalities of Inari, Kivijärvi, Kuhmo, Kuivaniemi, Kuusamo, Oulu, Pudasjärvi, Salla, Suomussalmi and Yli-Ii), mostly between 1973 and 1990. Detailed information of individual number, sex, finding date, municipality and additional data provided by the museum database (weight, body, tail and ear lengths, etc.) are available in the Supplementary Information #2.

Measurements were taken using an electronic calliper. Shreds of desiccated tendon sometimes present at the ends of bones were removed before taking measurements. To ensure replicable, usable measurements, we sought to make the measurement process as intuitive as possible, using the calliper in such a way that the gesture could easily be correctly reproduced (e.g., two contact points on the same face, allowing the calliper to be set correctly). From 9 to 10 measurements were taken for each bone, for a total of 55 measurements. All of the 55 measurements are illustrated in Figure 1, and descriptions are given in Supplementary Information #1.

**Figure 1:**
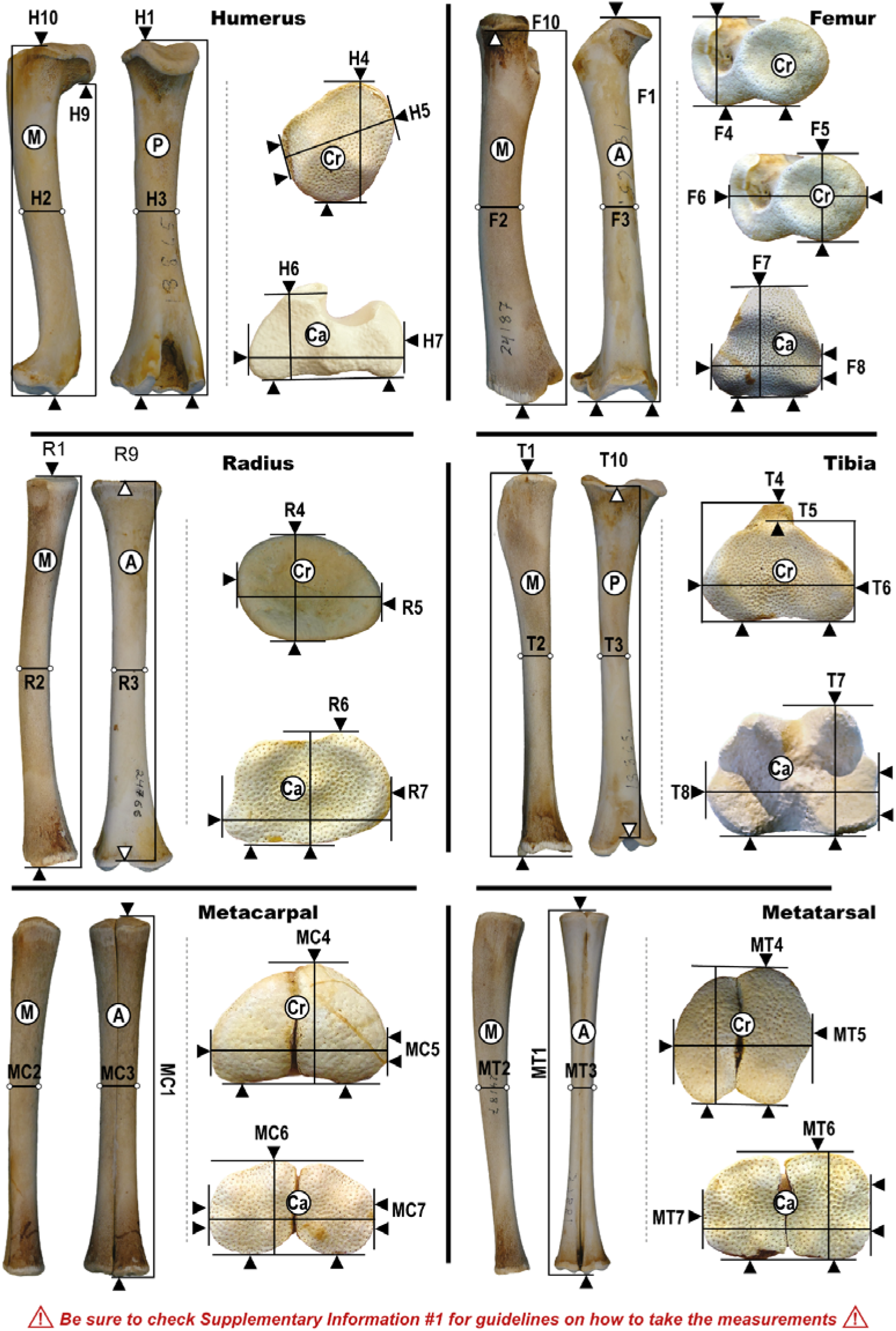
Illustration of calliper positioning for measurements of Rangifer foetal bones. Detailed descriptions and important guidelines are available in Supplementary Information #1.

The R8, H8, MC8, MT8, F9, T9 measurements correspond to the tightest point identified by sliding the calliper all along the diaphysis from top to down according to a medio-lateral axis. This point is roughly situated at mid-diaphysis but its location may vary slightly from one individual to another.

Statistical analysis and graphs were performed using R 4.2.2 (R Core Team, 2022) and RStudio (RStudio Team, 2020), with the packages this.path (Simmons, 2023), readxl (Wickham & Bryan, 2023), ggcorrplot (Kassambara, 2023), psych (Revelle, 2023), ggplot2 (Wickham, 2016), scales (Wickham, Pedersen, et al., 2023) and dplyr (Wickham, François, et al., 2023). The interface was designed using the shiny package (Chang et al., 2023). The R codes are available at the following website: https://github.com/ediscamps

## Results

To achieve the goals set in the introduction, a three-step approach was followed: 1) test whether the different measurements are correlated with each other, and thus if they can all be used to estimate one another as well as foetus overall size (i.e. which skeletal measurements are useable to assess foetus size? are long bones growing at a similar pace during foetal development?); 2) estimate how foetus skeletal measurements are correlated with foetal age; 3) conceive a digital interface for the estimation of foetal age from skeletal measurements of fossil bones.

### Correlation between skeletal measurements

Nearly all measurements (53 out of 55) are highly correlated between each other (Figure 2). H8, MC8 and MT8 (which are measurements taken at a variable point on the bone - the thinnest part of the diaphysis) have non-significant correlations with other measurements, 9 raising doubts on their interest as a measure of foetus overall size: they were excluded from further analyses. With these three measurements excluded, all correlations are significant (p < 0.05), and the Pearson r correlation statistic varies between 0.69 and 0.99 with a mean of 0.93, attesting to strong linear positive correlations. Figures 3, 4 and 5 present correlations and bi-plots between measurements in more detail. R1 stands out from others by showing a non-normal distribution (Figure 3), but this is mostly due to the fact that this measurement could only be taken on part of the specimens.

**Figure 2 :**
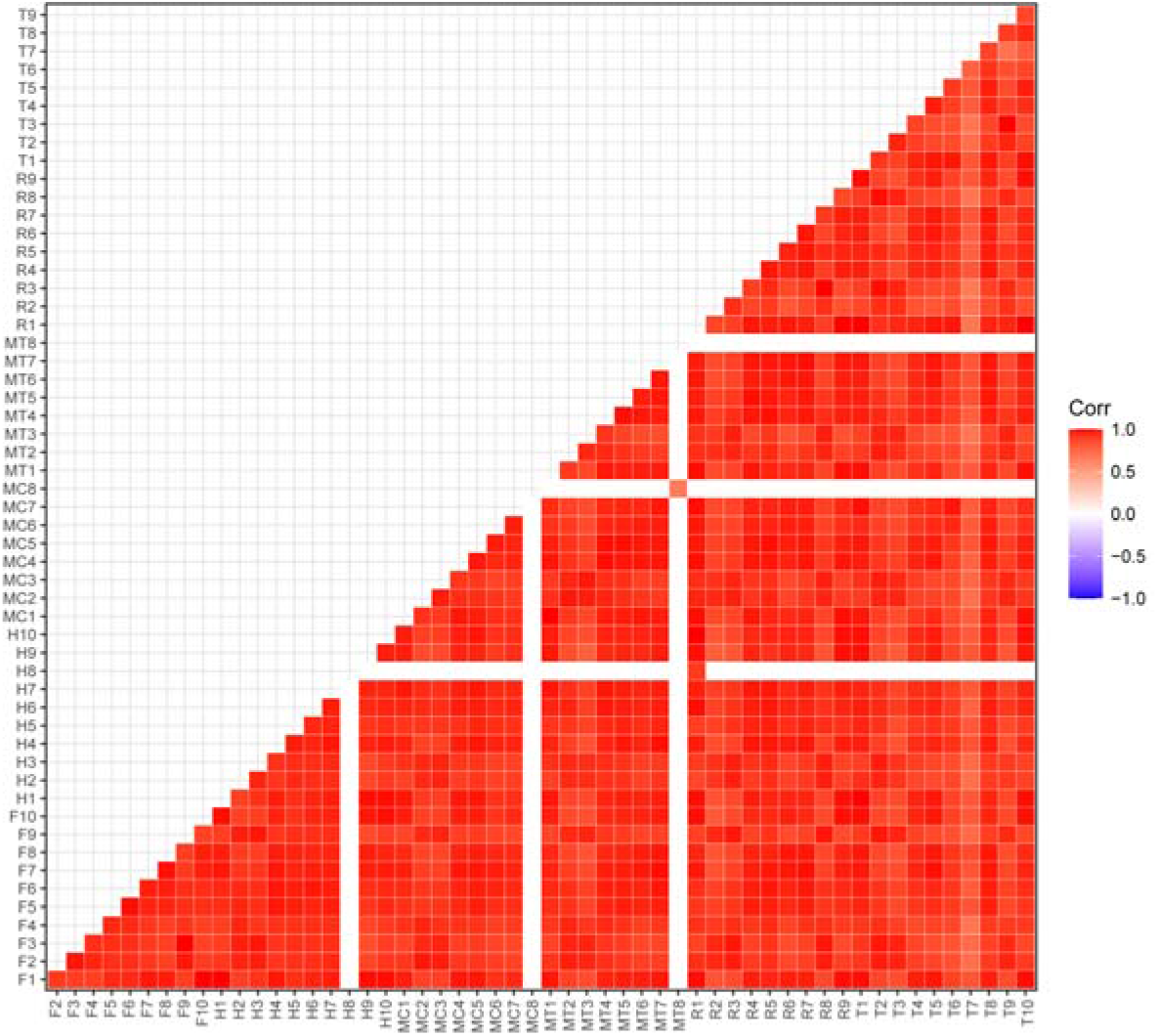
Pearson correlations between measurements. Statistically non-significant (p > 0.05) correlations are represented by blank cells.

**Figure 3 :**
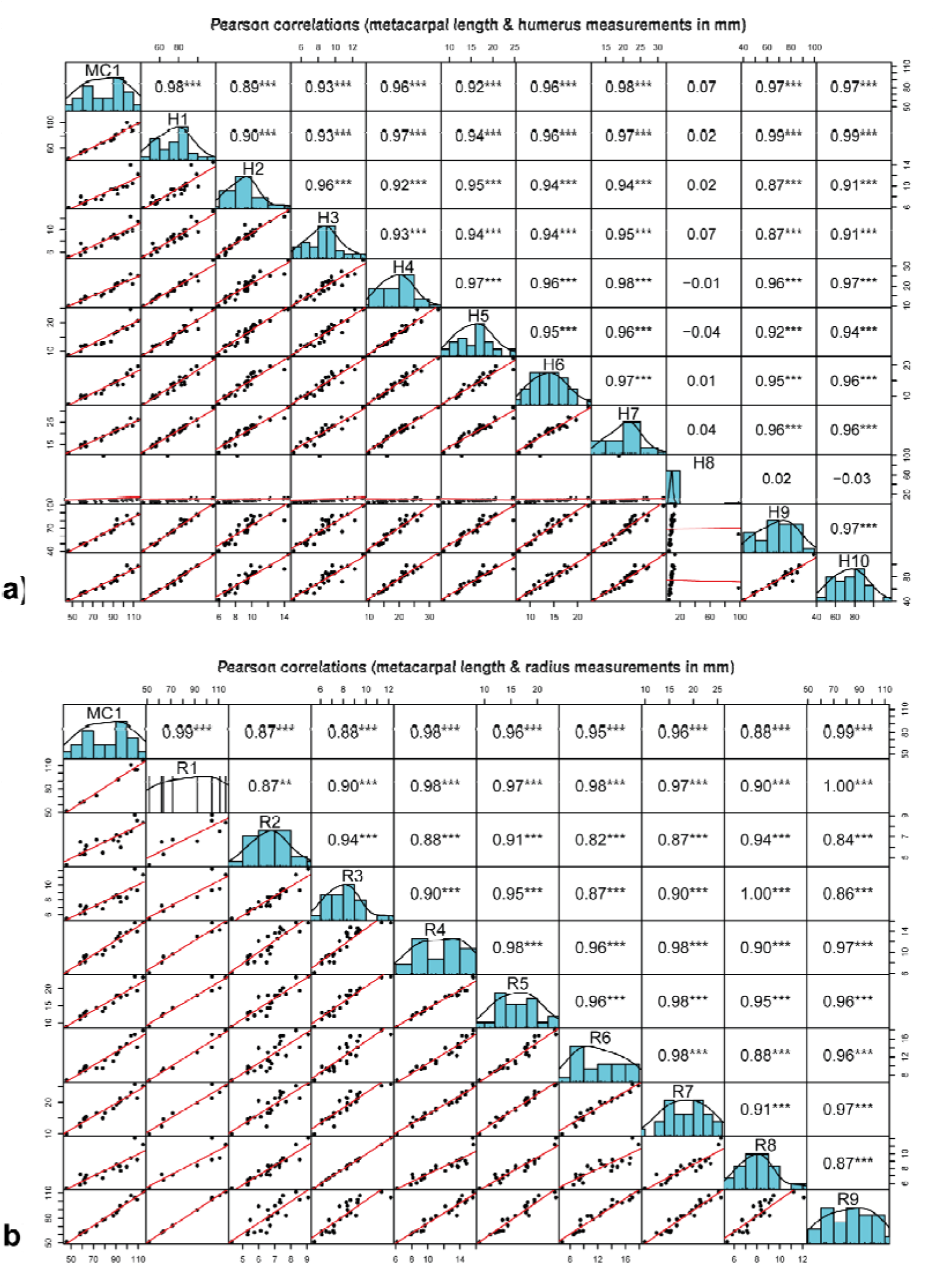
Humerus (a) and radius (b) measurements against metacarpal length. Diagonal:

**Figure 4 :**
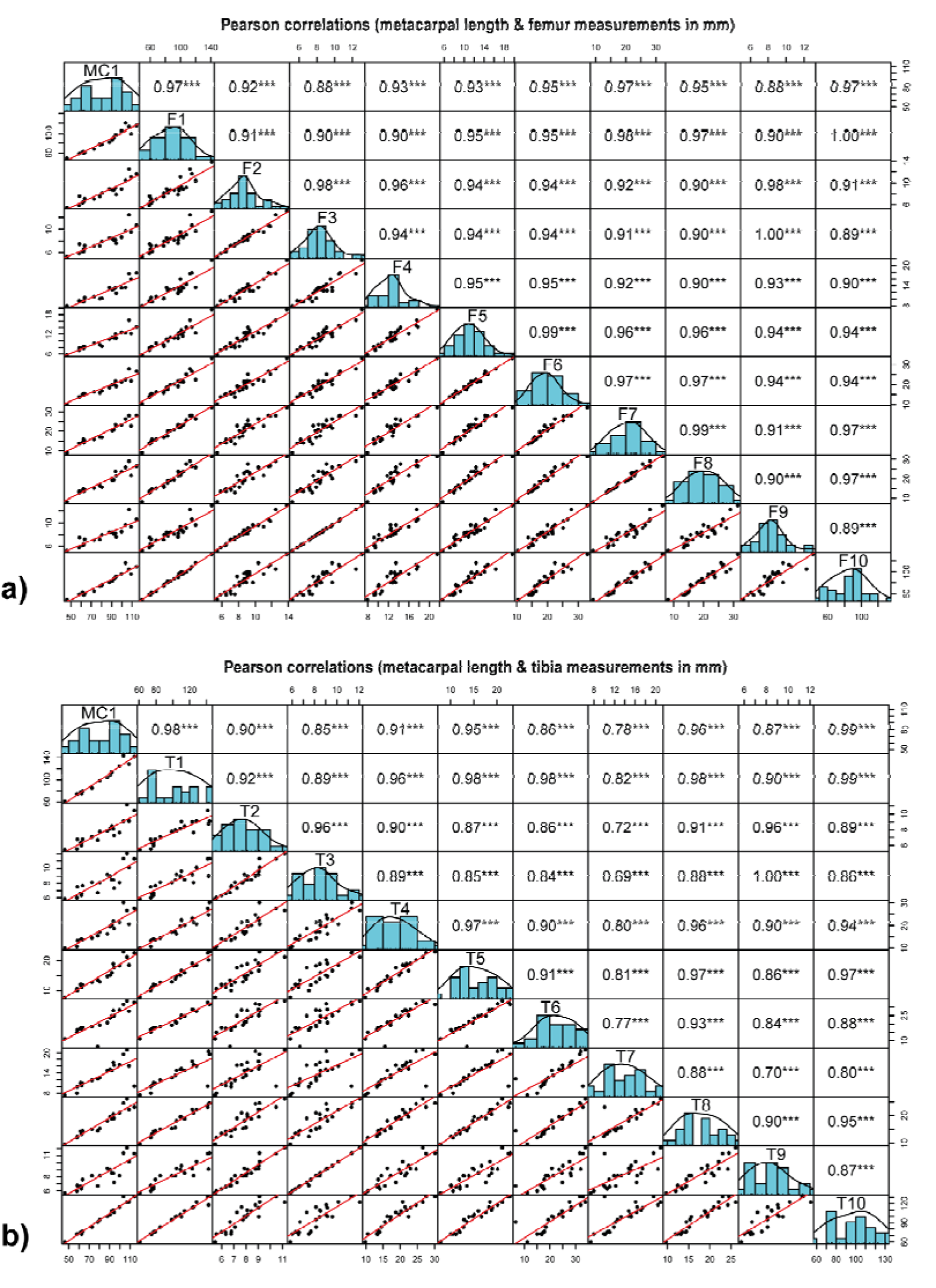
Femur (a) and tibia (b) measurements against metacarpal length. Diagonal:

**Figure 5 :**
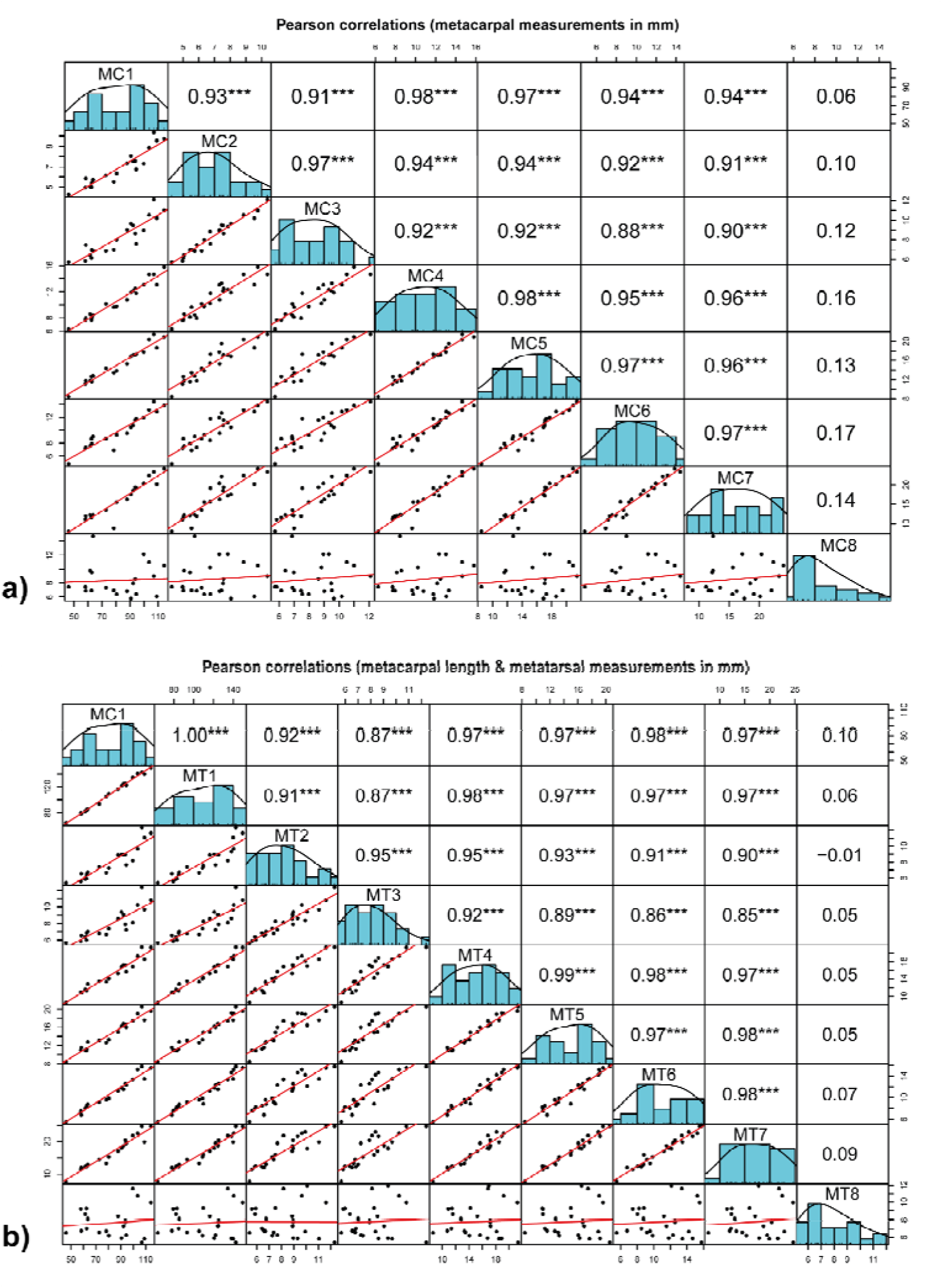
Metacarpal (a) and metatarsal (b) measurements against metacarpal length. Diagonal: histogram of measurements. Lower triangle: bivariate scatterplots of measurements. Upper triangle: Pearson correlation statistic (*: p < 0.05; **: p < 0.01; *** : p<0.001).

Overall these results show that all elements grow at a similar pace during foetal development. Apart from H8, MC8 and MT8, any measurement can be used to estimate the foetus overall size, as well as other measurements from the same bone, or from another one (e.g. for the establishment of Minimum Number of Individuals).

### Estimating the foetal age of modern individuals

Skeletal measurements should be strongly linked to foetal age, yet this variable is hardly ever known, even for modern individuals. Indeed, the exact foetal age depends on two events, conception date and birth, that are, especially for the former, not always known. Exploring links between foetal size and age are thus not straightforward.

Available data on rut and calving periods are summarised in Supplementary Information #3. In Finland, Norway (Svalbard excluded) and Sweden, rut periods reported by different authors span from the 25th of September to the 19th of October, and calving periods from the 5th of May to the 7th of June, with a peak around mid-May (SI #3). For reindeer, implantation is not delayed (Leader-Williams & Rosser, 1983), and Roine *et al*. (1982) considered that the majority of hinds had conceived by the 10th of October. Estimates of the gestation period vary between 209 and 240 days (Leader-Williams & Rosser, 1983), while a mean gestation period of 221 days (211 to 229) was reported for reindeer populations at the Kutuharju Field Reindeer Research Station, in Kaamanen, Finland (Mysterud et a.l, 2009), similar to the estimate of 220 days by Roine *et al*. (1982).

Thus, despite an important variability, we used in our study mean values for conception and birth around, respectively, the 10th of October and the 18th of May, with a mean of 221 days of gestation.

In our sample, the exact development age of each individual at death is only known for 7 individuals (Table 1). A finding date was recorded in the Oulu Museum database, but the exact time span between death and foetus discovery is, in most cases, unknown (however, given that they are very attractive to predators, who would have destroyed or at least damaged the bones, it is likely that they died shortly before being found). Rather than using this small sample (n = 7), we estimated foetal age (in weeks) using data provided by Roine et al. (1982), who studied a much larger sample also from Finland. Using their data (Roine et al., 1982: their Table 1), the following 2-degree polynomial regression equation between mean metacarpal length (MC1) and foetal age (in weeks) was obtained: y = -0.0015x^2^ + 0.3691x + 7.808 (R² = 0.9961). MC1 measurements from our sample were then used to estimate foetal age (in weeks). For 11 individuals, MC1 could not be measured: F2 was thus used to estimate MC1 (F2 and MC1 are highly correlated, r = 0.92 ; p < 0.001), and subsequently the estimated age. The comparison of estimated age with the known age of 7 individuals shows good agreement, with a mean difference of 2.44 weeks, and a maximum of 5.64 weeks (Table 1). These differences could be due to a different date of conception (here, all ages using the death date assume a conception on the 10th of October) or issues with the estimated age using skeletal measurements.

**Table 1:** Summary of information provided by the Oulu Museum on the death of seven individuals. 1: the “MeanAge” column corresponds to the number of weeks between a mean date of conception (10th of October) and the date of death. 2: “EstimAge” is the estimated age (in weeks) computed using skeletal measurements (cf. text for more details).

| Individual | Death information | Finding date | MeanAge<sup>1</sup><br>(weeks) | EstimAge<sup>2</sup><br>(weeks) | Difference<br>(1 - 2) |
| --- | --- | --- | --- | --- | --- |
| 18861 | in the uterus from bear-killed mother | 31/05/1979 | 33.43 | 27.79 | +5.64 |
| 18862 | in the uterus from wolverine-killed mother | 24/04/1979 | 28.14 | 28.93 | -0.79 |
| 18863 | in the uterus from wolverine-killed mother | 28/04/1979 | 28.71 | 29.09 | -0.37 |
| 18865 | 1 day old, killed by bear | 31/05/1979 | 33.43 | 29.06 | +4.36 |
| 18868 | 0 day old, died in labour | 28/05/1982 | 33.00 | 30.30 | +2.70 |
| 24273 | 4 days old | 24/05/1989 | 32.43 | 30.40 | +2.03 |
| 24766 | about 1 day old | 15/05/1990 | 31.14 | 29.94 | +1.20 |

Figure 6 compares finding date, estimated age and several independent measurements of foetal size (weight and body length as recorded in the Zoological Museum of the University of Oulu database, MC1 and F2 skeletal measurements). Estimated age and finding date are correlated (r = 0.88), but estimated age is more strongly correlated with the different measures of foetal size (Figure 6). Hence, the estimated age computed in this study appears to be a better estimation of foetal age compared to the finding date.

**Figure 6:**
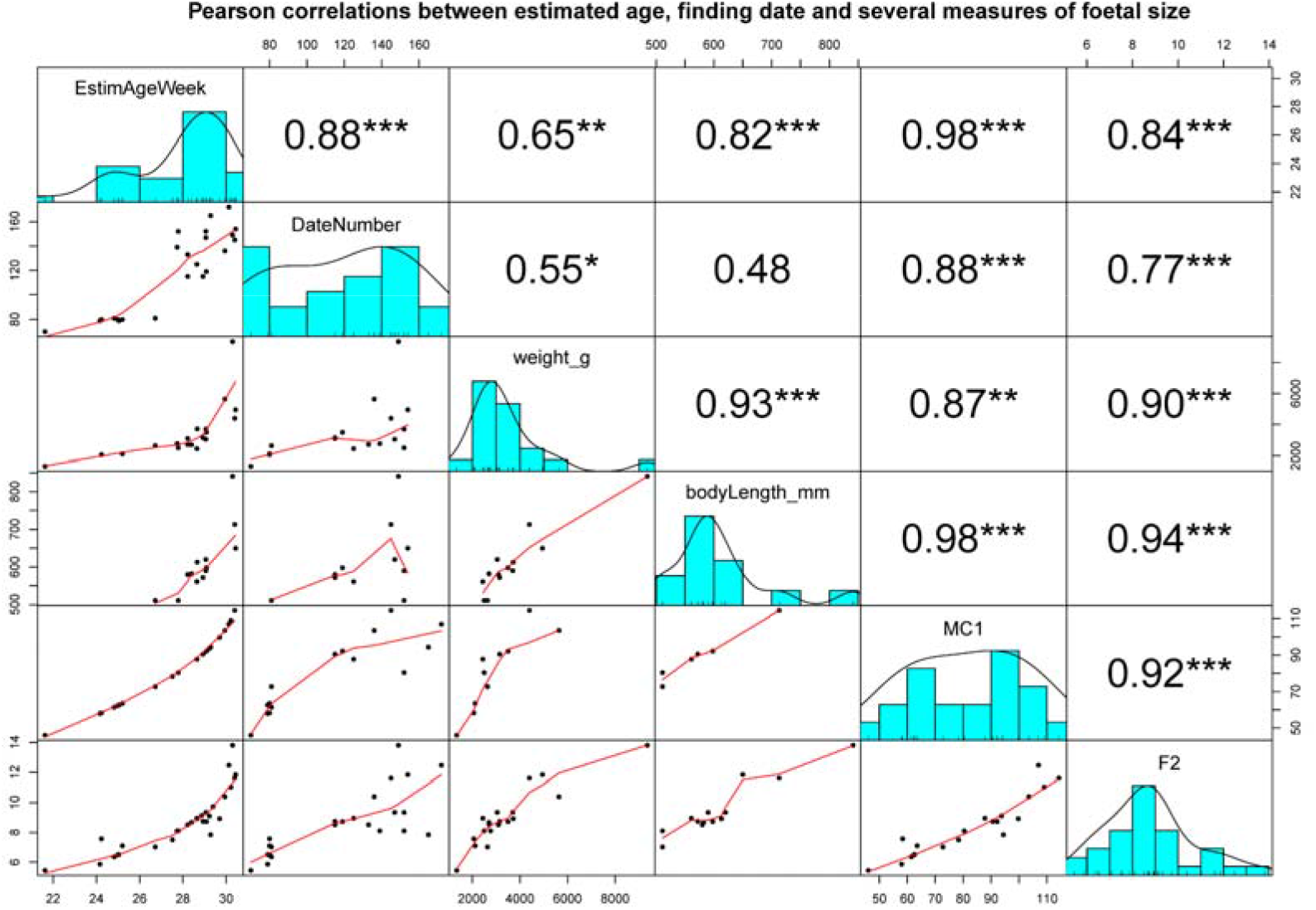
Correlations between estimated age (weeks, cf. main text), finding date (expressed as number of days since the 1st of January), foetal weight (g), body length (mm), metacarpal and humerus lengths (mm). Diagonal: histogram of measurements. Lower triangle: bivariate scatterplots of measurements. Upper triangle: Pearson correlation statistic (*: p < 0.05; **: p < 0.01; *** : p<0.001).

### Estimating foetal ages using skeletal measurements of fossil bones

Linear modelling with 2-degree polynomial regression equations were then computed between each skeletal measurement and the estimated foetal age (in weeks). Such equations allow for the estimation of foetal age with any skeletal measurement.

The high number of measurements (n = 55) renders the publication and use of regression curves highly impractical for zooarchaeologists. Thus, a R Shiny application called “foetusmeteR” was designed to allow for:

- The estimation of development age and season of death (Figure 7) using any of the measurements listed in Table 1 and Figure 1.
- The quick calculation of one measurement according to another (Figure 8), to facilitate the calculation of Minimum Number of Individuals (e.g. “can a femur of this size be attributed to a humerus of that size?”);

**Figure 7:**
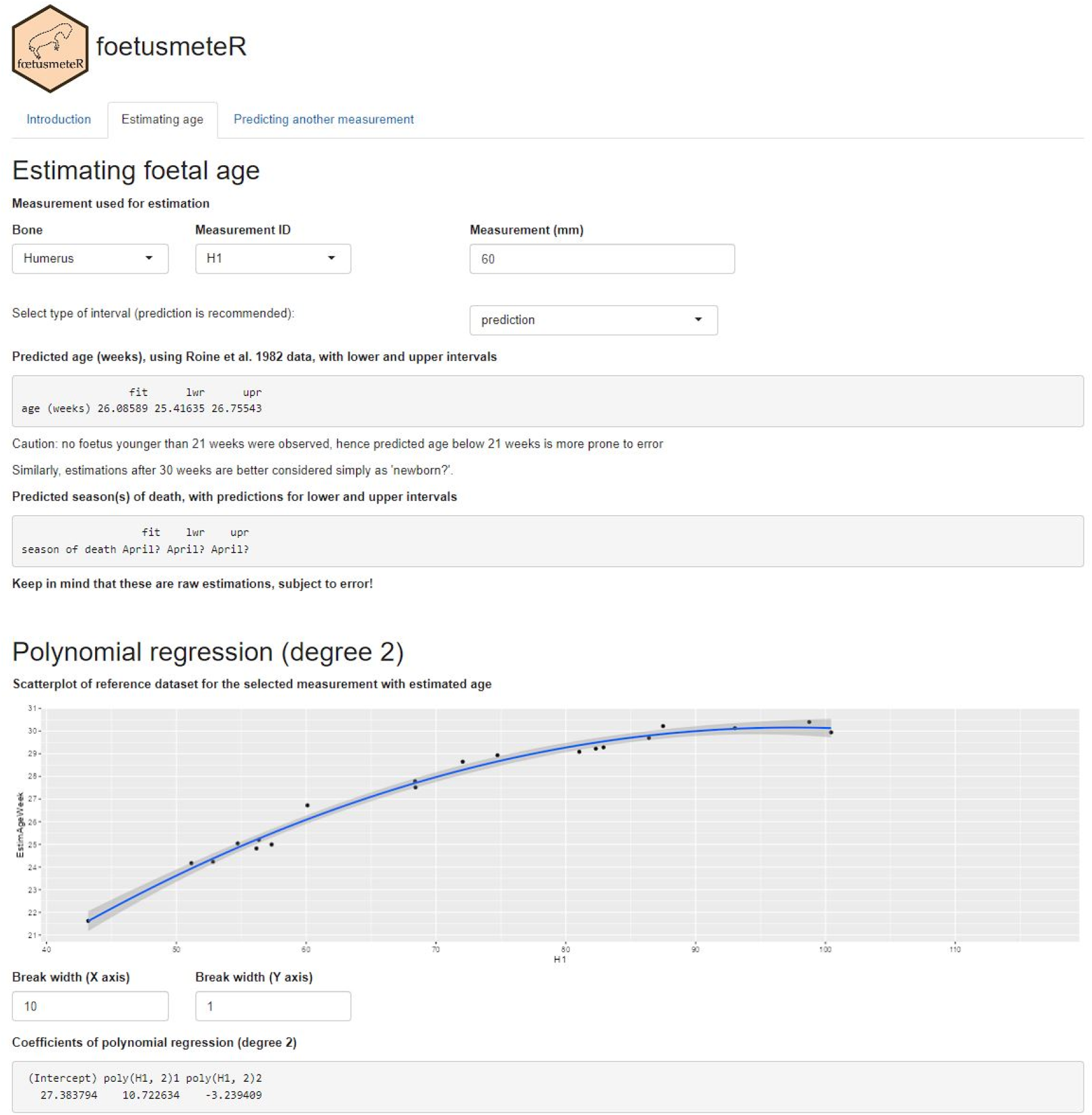
Screenshot of the first panel of the foetusmeteR interface that allows for the quick calculation of estimated foetal age and predicted season of death.

**Figure 8:**
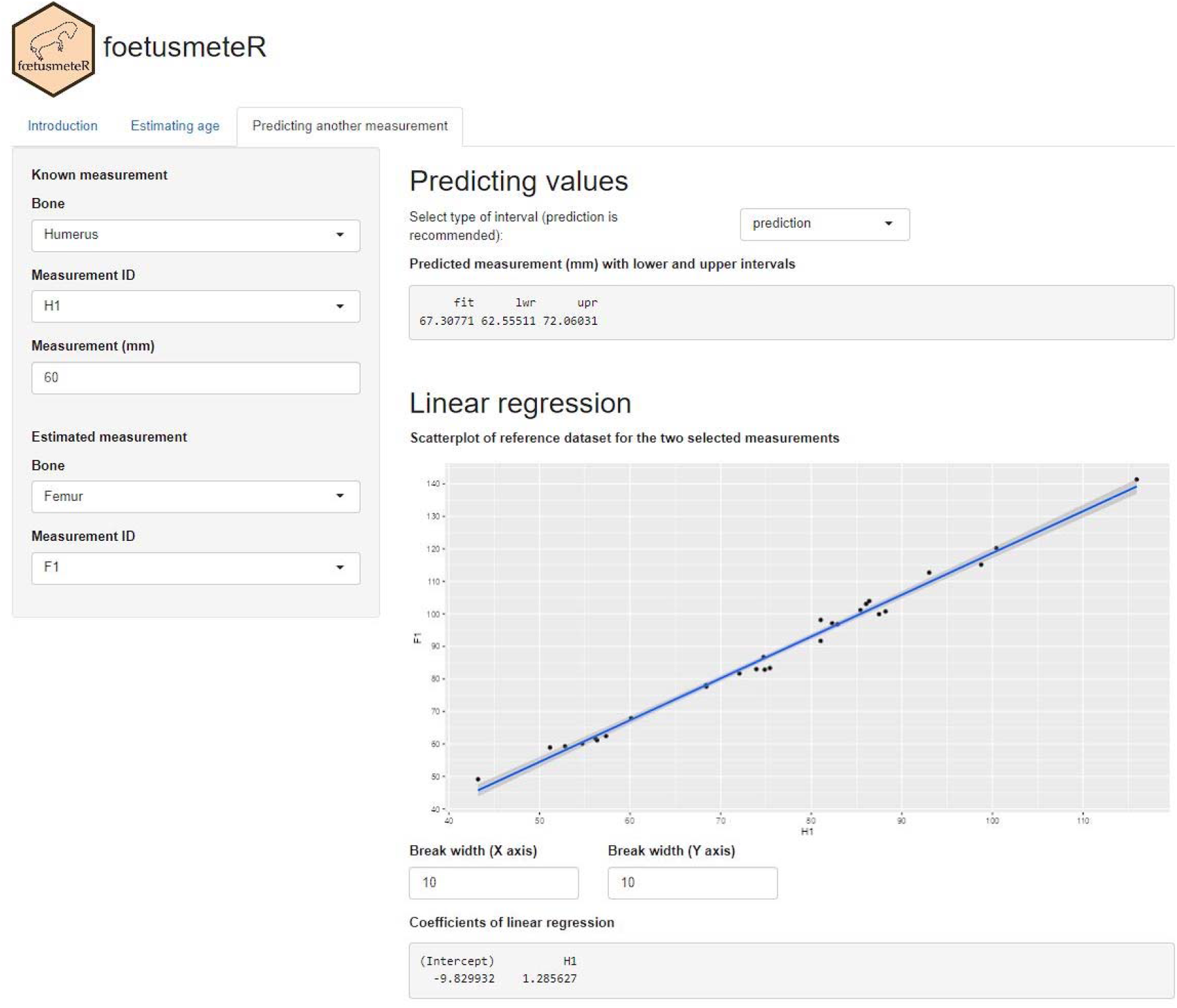
Screenshot of the second panel of the foetusmeteR interface that allows for the prediction of an unknown measurement using a known measurement.

The foetusmeteR interface is freely available for download at https://github.com/ediscamps/foetusmeteR, or for online use at https://ediscamps.shinyapps.io/foetusmeteR/

When only raw estimates are looked for, “cheat sheets” with scaled photographs are also made available in Figures 9 to 11: once printed, these pages can be used by simply placing the foetus bone on top of it for a quick and easy identification of season of death. Besides, for small fragments that cannot be measured, the cheat sheet can be used to estimate other measurements, giving access to the size of the complete foetal bone, and then to the age of the foetus.

**Figure 9:**
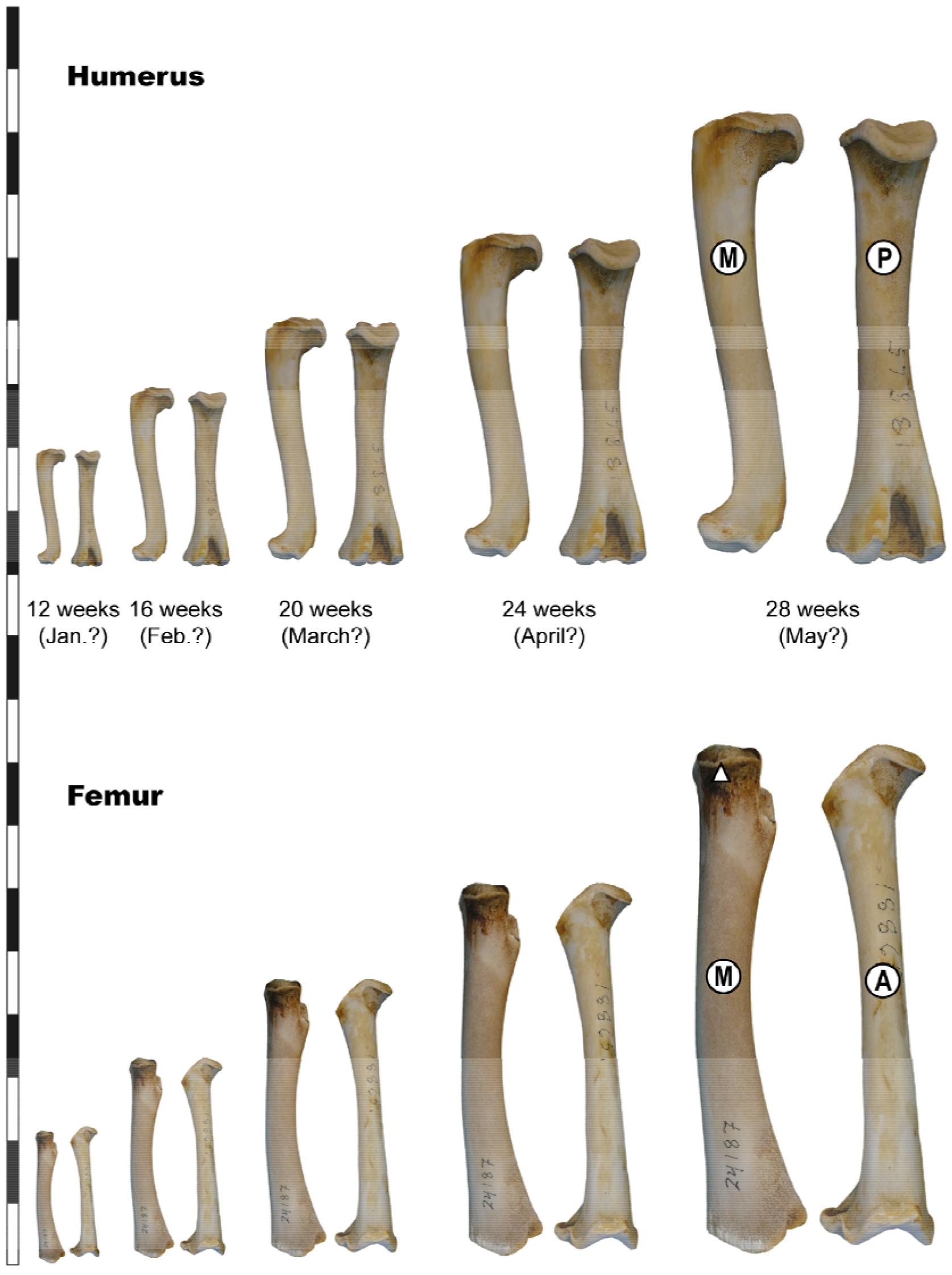
Foetus “cheat sheet” showing the average size for humerus and femur at different

**Figure 10:**
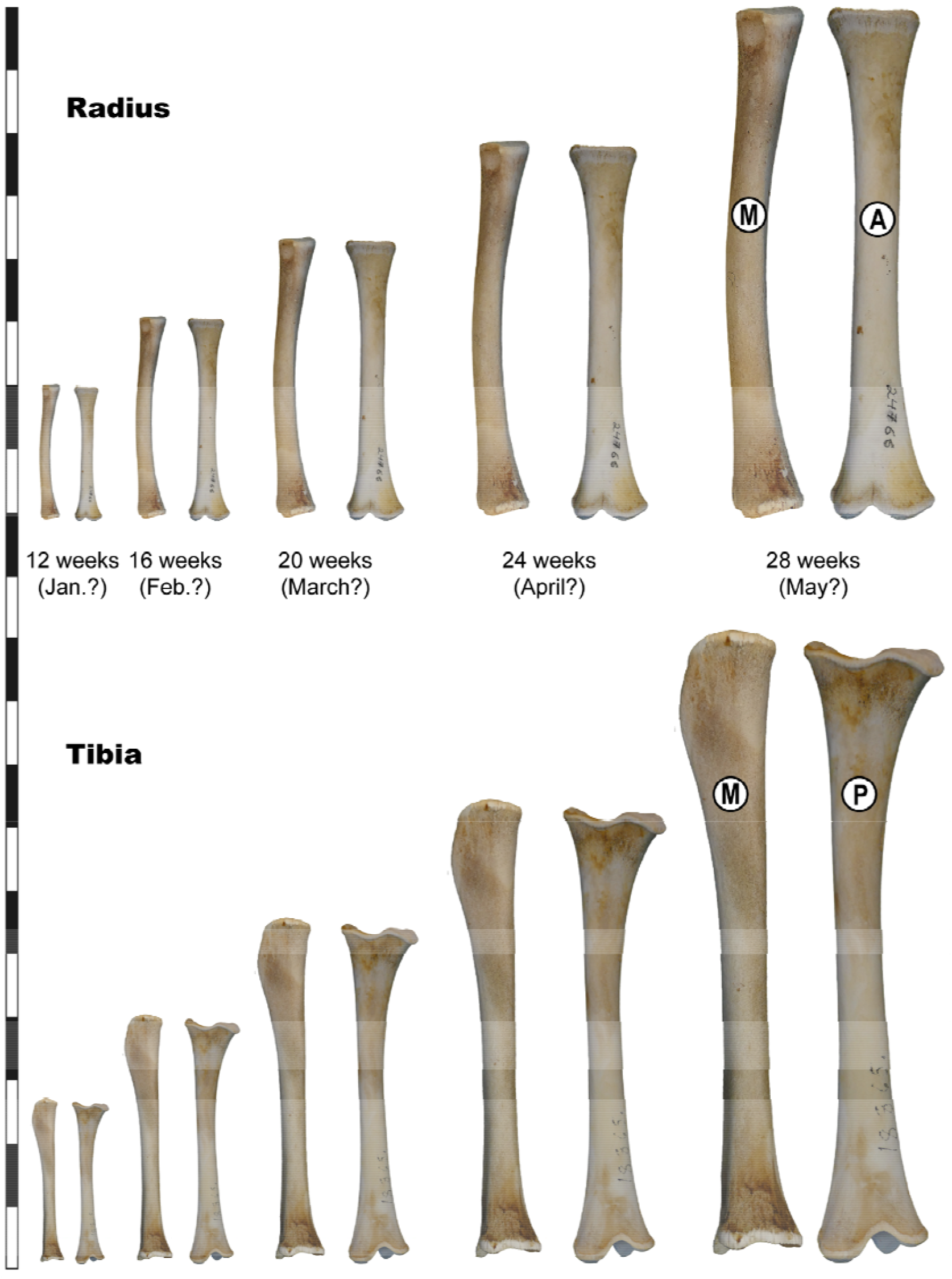
Foetus “cheat sheet” showing the average size for radius and tibia at different

**Figure 11:**
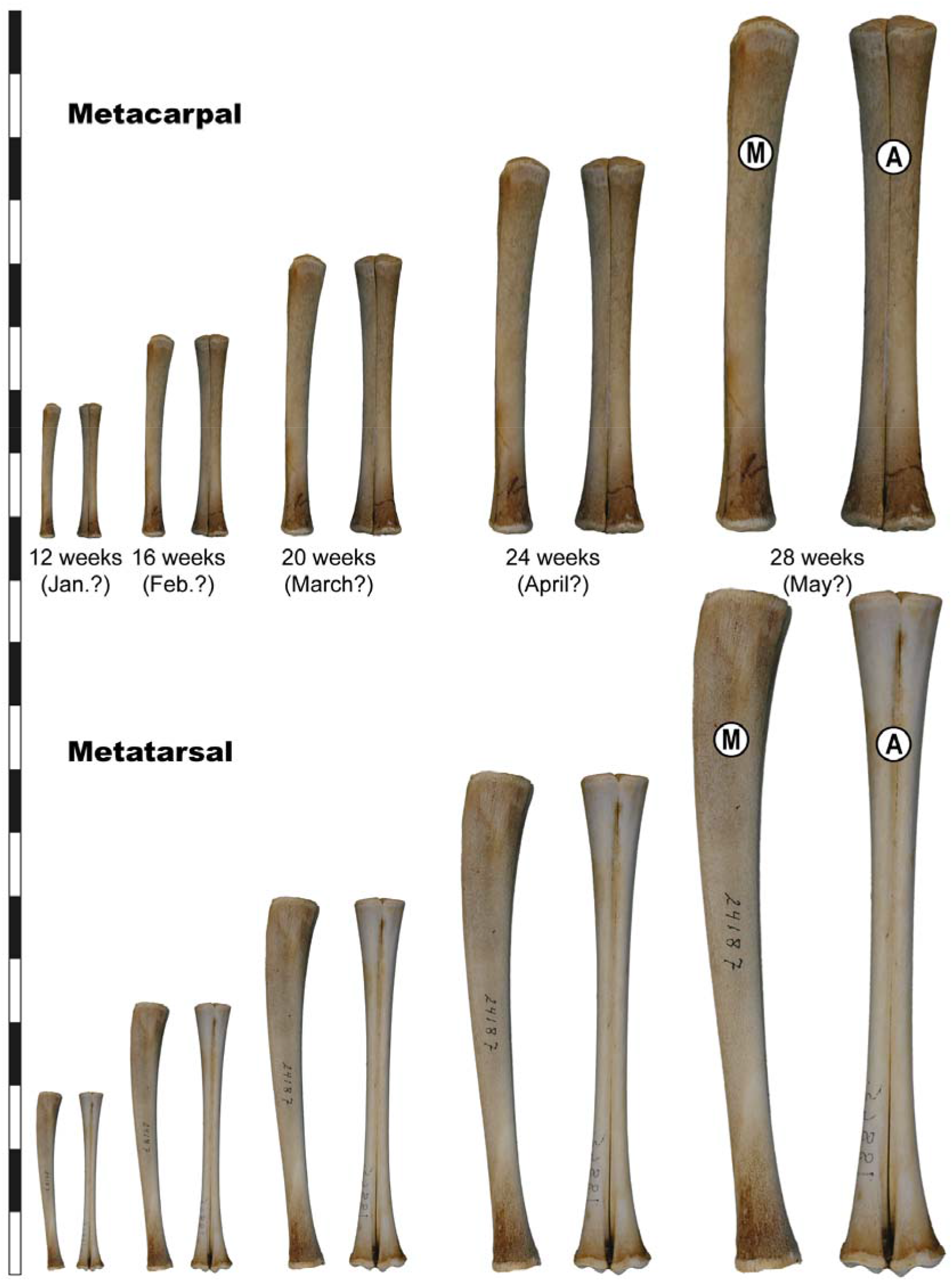
Foetus “cheat sheet” showing the average size for metacarpals and metatarsals at different foetal ages. Once printed at the correct scale, this cheat sheet can be used to quickly assess the age of a foetus bone placed on top of it.

## Discussion

Several biases affecting the method proposed here should be considered, pertaining mostly to known differences between populations, subspecies and sexes.

First and foremost, adult size is known to vary across *Rangifer* subspecies and sex (Geist, 1999). For example, adult wild forest *R.t. fennicus* are about 11-18% taller than semi-domesticated *R. t. tarandus* (Nieminen & Helle, 1980), and these differences are reflected in skeletal measurements (Puputti & Niskanen, 2008; Pelletier et al., 2020, 2022). The same can be said between males and females. To what extent these size differences are reflected in foetal size at a given age, or foetal growth speed, is hard to grasp with the datasets at hand. Mean birth weights of Finnish *R.t. tarandus* were reported to be slightly larger for males (5.3kg) compared to females (5.0kg), thus attesting to a 6% difference at the end of foetal growth (Nieminen & Petersson, 1990).

In our dataset, for *Rangifer tarandus tarandus*, regression curves obtained for males (n = 11) or females (n = 11) are extremely similar, pointing to a near-zero influence of sex (Figure 12a). When the dataset is split by subspecies, regression curves obtained for *Rangifer tarandus fennicus* (n = 6) are slightly offset from the ones obtained for *Rangifer tarandus tarandus*, yet confidence intervals overlap (Figure 12b). In the absence of a larger sample of *R. t. fennicus* foetus of known date of death, the influence of subspecies on foetal growth can be considered as minimal for the intended goal of the study (i.e. broadly estimating foetal age), even though *R.t. fennicus* would be expected to be slightly larger.

**Figure 12:**
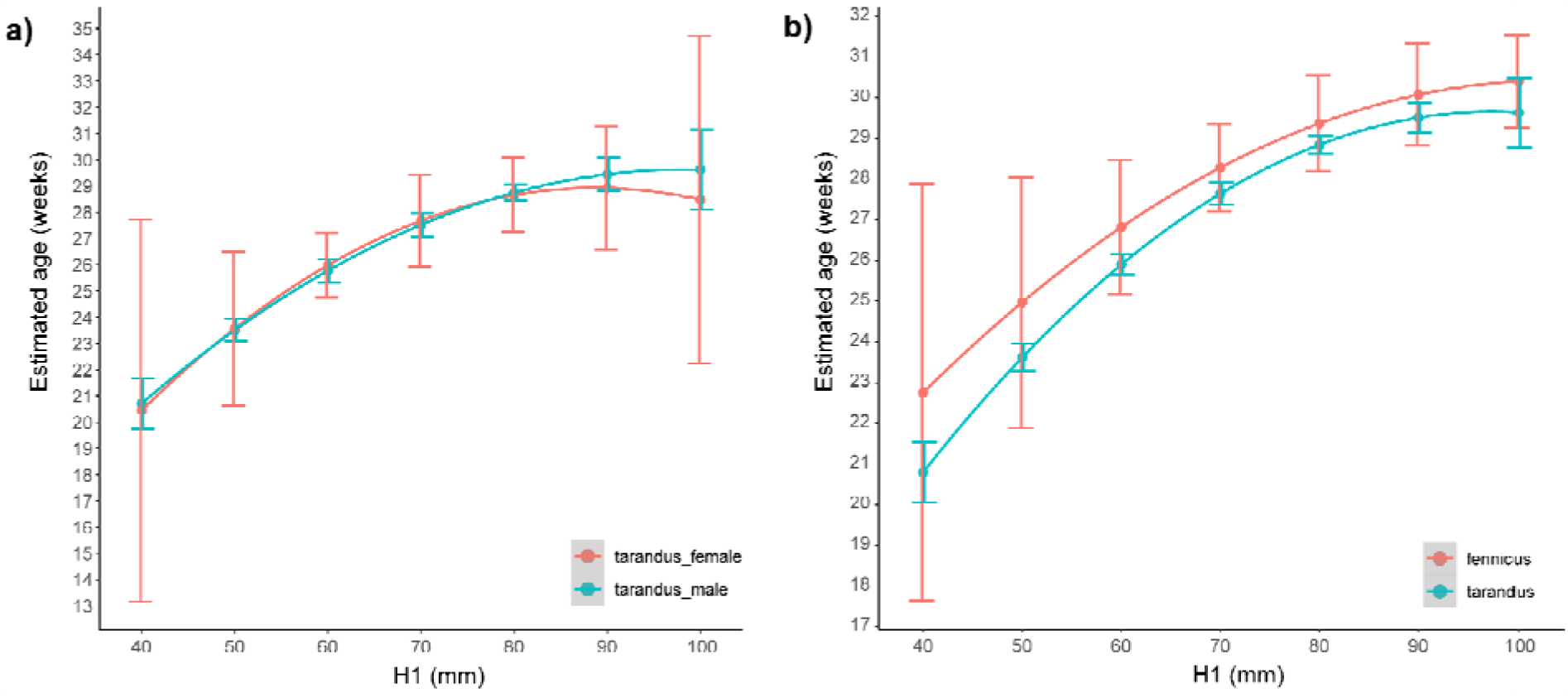
Differences between regression curves (with confidence intervals) obtained between H1 skeletal measurement (mm) and estimated age (weeks), when the dataset is separated by sex (a, for R.t. tarandus) or by subspecies (b). Similar results were obtained for all skeletal measurements.

Finally, one should keep in mind that, even if we assume that foetal age could be broadly assessed for all subspecies and sexes using the same equations, there is still a variability in rutting and calving periods across populations and subspecies (Supplementary Information #3). Delayed conception and calving are hypothesised to be a consequence of relatively poor physical condition of females in autumn (Reimers, 1983a; b; Flydal & Reimers, 2002). Thus, even if the foetal age is correctly predicted through skeletal measurements, the seasonality assessed from it should be regarded as an estimate.

In Finland, Norway and Sweden, rutting and calving periods are centred around, respectively, mid-October and mid-May. Rutting periods are overall similar for caribou populations with the following reported spans (Supplementary Information #3): early October to mid-November for *R. t. caribou* (Canada), 6th to 23th October with a peak around the 15th of October for *R. t. granti* (Alaska), and 19th to 31st of October for *R. t. groenlandicus* (Alaska, Canada). Calving is, however, more variable, especially at higher latitudes (Gunn & Fournier, 2000). For *R. t. caribou* (Canada), calving periods were reported from the 1st of May to 25th of June. For *R. t. granti* (Alaska), births occur from the 17th of May to the 13th of June, with a peak around the end of May. For *R. t. groenlandicus*, calving seems more centred around the third week of May (13th-21th) in Alaska, while, in Canada, births occur later, in June (8^th^ - 15th). Accordingly, gestation (around 221 days for *R. t. tarandus*) is longer for free-ranging North American caribou (around 225 to 235 days) (Skoog, 1968; Miller, 2002). Such differences should be kept in mind when using foetusmeteR in North American contexts.

All in all, all these biases and limits call for caution. Pending more work on these issues (e.g. inter-population variability, sexual dimorphism), foetal age and season of death are best estimated with the method proposed here, but such figures should only be used with full knowledge of their inherent limits, especially when applied to fossil assemblages.

## Conclusion

In the archaeological record, seasonal data is mostly obtained either by the observation of eruption and wear sequences of deciduous teeth (a method limited to very young individuals), or by carrying out cementochronological studies (a destructive method based on the observation of cementum increments along adult tooth roots). These two methods often provide relatively low resolution seasonal information, with estimations of periods of death extending over several months. The measurement of foetus bones is another method that can give fairly accurate results, and it allows the seasonality of hunting to be assessed on the basis of a different proxy (i.e. skeletal measurements) based on another sub-population of reindeer herds (i.e. pregnant females). Combining seasonal data acquired from different proxies and sub-populations can give a better picture of the hunted groups.

According to Spiess (1979: p. 187), bones of a foetus of less than 60 days are unlikely to be preserved in archaeological sites. Thus, the presence of foetus bones can correspond to hunts between December and May, winter and spring: without measurements, the seasonal information accessible to zooarchaeologists is broad and imprecise (e.g. Castel, 1999; Bemilli & Bayle, 2009; Connet et al., 2012; Pétillon et al., 2015; Castel et al., 2017; Dachary et al., 2020) and impedes discussion about hunting strategies and techniques. Indeed, during this period of time, the sanitary condition of the reindeer undergoes major changes (e.g. Spiess, 1979; Finstad & Prichard, 2000; Soppela & Nieminen, 2001) and, when hunting is your only means of subsistence, this detail is far from insignificant. Similarly, it is not the same in terms of hunting strategies and techniques to track isolated reindeer in covered areas (in winter?) as it is to plan an interception of a migrating herd (in spring?). The results of our study on foetal bones, applying a larger set of measurements on all long bones, allow for a finer estimation of prey acquisition seasonality on a wider array of archaeological contexts (e.g. even when foetus bones are fragmented). In the end, such renewed data should contribute to a better understanding of the nomadic lives of past human groups interacting with reindeer herds.

## Acknowledgments

This research was funded by the DeerPal project (Humans and deer during the Palaeolithic: integrating the variability of prey ecology and ethology in the investigation of past human— environment interactions; French National Research Agency: ANR-18-CE03-0007). Our deepest thanks go to the Zoological Museum of the University of Oulu and its members, and especially to Tuula Pudas for her precious help in accessing the material studied here, and for all the information provided on the specimens. We also wish to thank Marie Matu and Maxime Pelletier for their help and warm hospitality.

## Supplementary Information

Supplementary Information #1: Description of measurements by bone, in relation to the codes illustrated in Figure 1. The last column gives important guidelines for certain measurements on how to hold the bone and/or the calliper.

Supplementary Information #2: Detailed information on the 31 foetus measured provided by the Zoological Museum of the University of Oulu, courtesy of Tuula Pudas (species, sex, finding date, municipality, measurements, etc.).

Supplementary Information #3: rut (conception) and calving (birth) periods for different *Rangifer* populations.

